# Pharmacological perturbation of intracellular dynamics as a SARS-CoV-2 antiviral strategy

**DOI:** 10.1101/2021.09.10.459410

**Authors:** William Bakhache, Emma Partiot, Vincent Lucansky, Yonis Bare, Boris Bonaventure, Caroline Goujon, Cédric Bories, Maika S. Deffieu, Raphael Gaudin

**Author notes:** Corresponding author: Raphael Gaudin, 1919 route de Mende, 34293 Montpellier, France.

## Abstract

SARS-CoV-2 (CoV2) is the viral agent responsible for the pandemic of the coronavirus disease 2019 (COVID-19). Vaccines are being deployed all over the world with good efficacy, but there is no approved antiviral treatment to date. This is particularly needed since the emergence of variants and the potential immune escape may prolong pandemic spreading of the infection for much longer than anticipated. Here, we developed a series of small molecules and identified RG10 as a potent antiviral compound against SARS-CoV-2 in cell lines and human airway epithelia (HAE). RG10 localizes to endoplasmic reticulum (ER) membranes, perturbing ER morphology and inducing ER stress. Yet, RG10 does not associate with SARS-CoV-2 replication sites although preventing virus replication. To further investigate the antiviral properties of our compound, we developed fluorescent SARS-CoV-2 viral particles allowing us to track virus arrival to ER membranes. Live cell imaging of replication-competent virus infection revealed that RG10 stalls the intracellular virus-ER dynamics. Finally, we synthesized RG10b, a stable version of RG10, that showed increased potency *in vitro* and in HAE with a pharmacokinetic half-life greater than 2 h. Together, our work reports on a novel fluorescent virus model and innovative antiviral strategy consisting of the perturbation of ER/virus dynamics, highlighting the promising antiviral properties of RG10 and RG10b.

## Introduction

Since the beginning of the COVID-19 pandemic, thousands of clinical trials of molecules and experimental therapies have been started worldwide. As of today, several vaccines based on the induction of anti-Spike antibody generation have been approved, giving some hope to shut-down this pandemic. However, preparedness is key, and because we are observing the emergence of variants that escape vaccination, complementary tools, such as antiviral development, remain highly needed.

Drug repurposing has been started at the onset of the pandemic, but this approach has not been successful to date. One strong candidate for antiviral COVID-19 treatment is Remdesivir, a nucleoside analog inhibiting SARS-CoV-2 *in vitro* and in preliminary clinical reports. However, it does not improve the condition of severe COVID-19 patients (Goldman et al., 2020), exhibits notable side-effects, and is not accessible to non-hospitalized patients because of the intravenous route of administration. Although further clinical investigations are still needed, one can anticipate that a direct-acting agent (DAA) targeting the viral polymerase, may not be sufficient to fight the disease in the long run, as RNA viruses tend to rapidly evolve powerful escape strategies through punctual mutations. As such, host-targeting agents (HTA) should also be developed. Specifically, the need for novel antiviral strategies is still highly required because 1) drug repurposing may fail or need chemical optimization to increase efficiency against SARS-CoV-2, 2) side effects may be too important to treat vulnerable populations, 3) viral mutation/adaptation could render potent antivirals less efficient, and 4) multitherapy is an interesting strategy to prevent/limit viral escape.

The Coronaviridae family is subdivided into four genera based on phylogenetic clustering: *Alpha-, Beta-, Gamma*- and *Delta-coronavirus*. SARS-CoV, SARS-CoV-2 (CoV2), and MERS-CoV cluster into the *beta* genus, while the human Coronavirus 229E (HCoV-229E) and NL63, and the feline coronavirus (FCoV) are from the *alpha* genus. The main outcome of CoV2 infection is acute respiratory distress syndrome (85%), but other clinical disorders exist, including acute cardiac and kidney injuries (31% and 23% respectively), secondary infection (31%), and shock (23%) (Huang et al., 2020). CoV2 is an enveloped positive-strand RNA virus exhibiting a ≈ 30 kb genome enwrapped into a particle composed of a cellular lipid bilayer and four structural proteins: Spike (S), Nucleocapsid (N), Envelope **(E)**, and Membrane (M).

CoV2 entry is highly dependent on the angiotensin-converting enzyme 2 (ACE2), similarly to its close relative SARS-CoV (Hoffmann et al., 2020), and shows partial dependency for TMPRSS2 (Walls et al., 2020). The virus is internalized in a clathrin-dependent manner and endosomal acidification is required if the target cells do not express TMPRSS2 (Bayati et al., 2021, Ou et al., 2021). Upon fusion with cellular membranes, the RNA genome of CoV2, protected by N, is targeted to ER membranes where translation of replication proteins occurs (V’Kovski et al., 2021, Wolff et al., 2020).

All the early stages of CoV2 infection are fully dependent on a variety of cellular machineries, and a better understanding of these steps is critical to devise novel antiviral strategies. Here, we present novel promising anti-CoV2 molecules and we explore the molecular mechanism conferring its antiviral activity.

## Results

### Identification of RG10 as a potent SARS-CoV-2 antiviral molecule

We had synthesized a series of sulfamoyl-phenyl derived compounds designed as potential antiviral agents. While most of them did not prevent CoV2 replication at 2 and 10 µM, we identified the molecule RG10, a novel small molecule of unknown target, as a potent antiviral molecule with similar activity levels as Remdesivir (Rem; Figure 1A). RG10 did not show cytotoxicity at working concentrations (Figure 1B) and measurement of its potency in Huh7.5.1 cells returned an EC_50_ ≈ 1.5 µM (Figure 1C).

**Figure 1.**
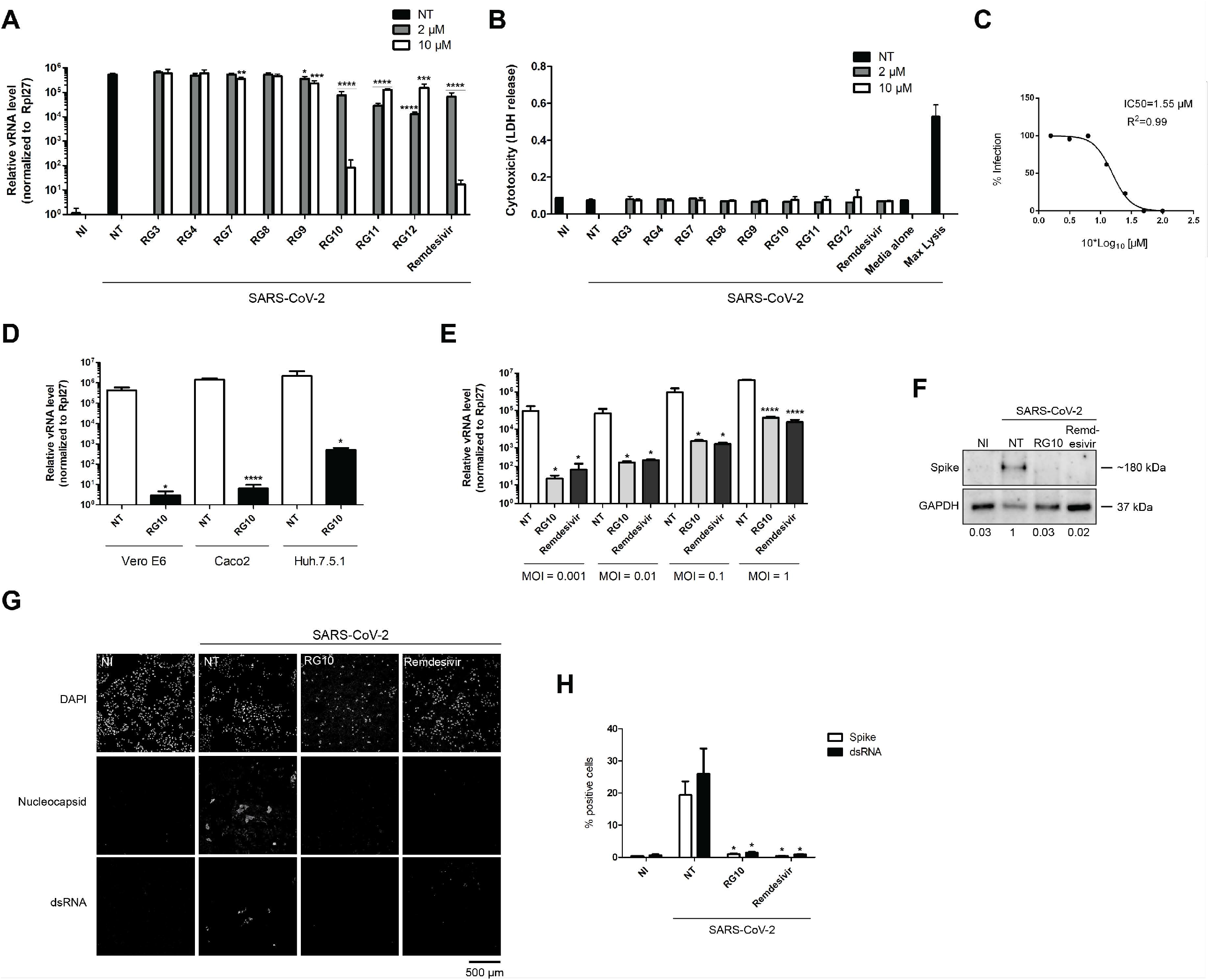
Identification of RG10 as a potent SARS-CoV-2 antiviral. **(A-B)** Vero E6 cells were treated with the indicated RG derivatives or Remdesivir at indicated concentrations and subsequently infected with SARS-CoV-2 at MOI 0.01 for 48 h in the presence of the compounds. **(A)** The graph shows relative SARS-CoV-2 RNA (vRNA) levels measured by RT-qPCR and normalized by Rpl27 RNA expression. The bars are means +/- SD from duplicates representative of two independent experiments. Student’s t-test ** p value < 0.01; *** p value < 0.005; **** p value < 0.001. **(B)** Cytotoxicity measured by LDH release assay showed no significant differences between the compounds. **(C)** Vero E6 cells were treated with RG10 at various concentrations and subsequently infected and processed as in **A**. The IC_50_ was 1.55 µM +/- 0.11 as measured from three independent experiments. **(D)** Vero E6 (monkey kidney cells), Caco2 (human intestinal cells) or Huh7.5.1 cells (human liver cells), were treated with 10 µM RG10 and subsequently infected with CoV2 at MOI 0.01 for 48 h in the presence of the compounds. The graph shows relative CoV2 RNA (vRNA) levels measured by RT-qPCR and normalized by Rpl27 RNA expression. The bars are means +/- SD from duplicates representative of two independent experiments. Student’s t-test ** p value < 0.01; *** p value < 0.005; **** p value < 0.001. **(E)** Huh7.5.1 cells were treated with 10 µM RG10 or 10 µM Remdesivir and subsequently infected with CoV2 at indicated MOI for 48 h. The graph shows relative vRNA levels measured by RT-qPCR and normalized by Rpl27 RNA expression. The bars are means +/- SD from duplicates representative of two independent experiments. Student’s t-test * p value < 0.05 and **** p value < 0.001. **(F)** Huh7.5.1 cells were treated with 10 µM RG10 or 10 µM Remdesivir and subsequently infected with SARS-CoV-2 at MOI 0.5 for 48 h. Cells were then lysed 48 h post infection. The lysates were processed for western blotting using an anti SARS-CoV-2 Spike antibody and an anti-GAPDH antibody as loading control. Both immunostainings were revealed using secondary antibodies coupled to HRP. (**G**) Cells were infected as in **F** then fixed and stained using an anti SARS-CoV-2 Nucleocapsid antibody and a J2 antibody recognizing double stranded RNA (dsRNA) representative of viral replication complexes. The micrographs show the individual channels that have been acquired using a confocal microscope and processed using ImageJ. (**H**) Samples were treated as in **F** and cells were detached and fixed 48 h post infection. Samples were stained and analyzed by flow cytometry. Student’s t-test * p value < 0.05.

RG10 exhibited high CoV2 antiviral activity in Vero E6 cells but also in human Caco2 and Huh7.5.1 cells (Figure 1D) and retained potency at high multiplicity of infection (MOI) similar to Rem (Figure 1E). By Western blot, neosynthesized Spike protein (S) could barely be detected upon RG10 or Rem treatment (Figure 1F). Microscopy and flow cytometry analyses confirmed that both RG10 and Rem prevent viral protein synthesis (Figure 1G-H). Moreover, immunolabeling using an antibody recognizing double stranded RNA (dsRNA), specific of viral replication complexes, showed that RG10 prevents viral RNA synthesis (Figure 1G-H).

### RG10 inhibits CoV2 replication

To discriminate whether RG10 inhibits virus entry or post-entry events, we performed time-of-addition experiments where the molecule was added for four hours and then washed-out (Pre), or conversely, no drug was added for the first 4 h to allow early virus entry steps to occur and only then RG10 was added to the cells (Post; Figure 2A). We found that RG10 had no antiviral activity when incubated only during virus entry, while adding RG10 post-entry significantly decreased vRNA expression levels (Figure 2B). In these conditions, comparing the amount of viral N protein and dsRNA levels by flow cytometry confirmed that RG10 exerts strong antiviral activity post entry, although some inhibition was also observed at the entry steps (Figure 2C-D). These results indicate that RG10 inhibits CoV2 replication, while inhibition of virus entry is likely minimal.

**Figure 2.**
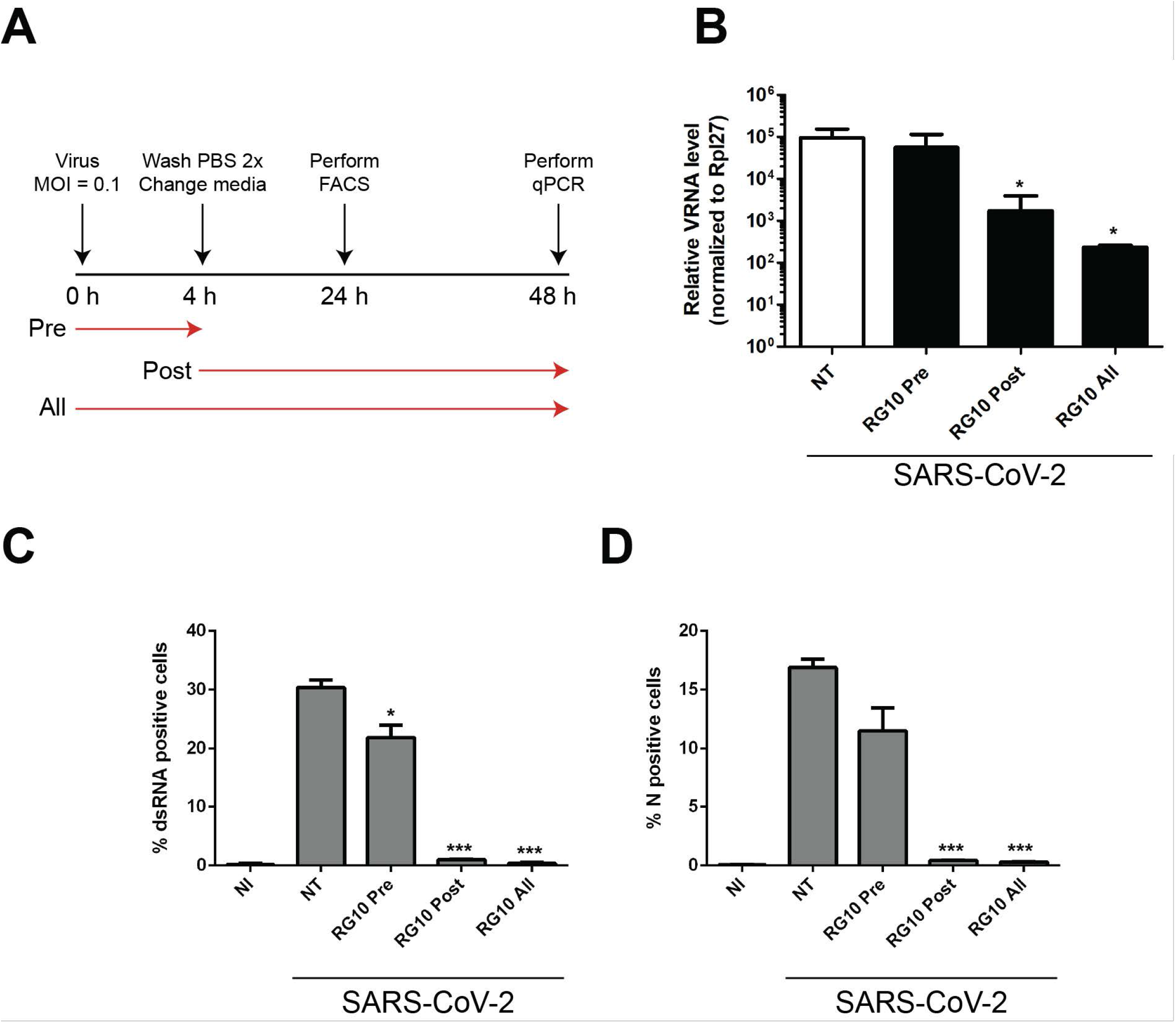
RG10 inhibits SARS-CoV-2 infection at a post-entry stage. **(A-D)** Vero E6 cells were non-infected (NI), or infected with CoV2 at MOI 0.1 (SARS-CoV-2). Cells were treated with 10 µM RG10 during the first 4 h of infection and washed-out (RG10 Pre), starting at 4 h post-infection (RG10 Post), or during the whole infection (RG10 all). A scheme representing the time-of-addition experiment is shown in **A**. Cells were lysed after 48 h and processed for RT-qPCR **(B)** or lysed after 24 h and processed for flow cytometry, staining for dsRNA and N protein **(C-D)**. The bar graphs are mean +/- SD from duplicates and representative of two experiments. Student’s t-test *** p value < 0.005.

### RG10 localizes to the endoplasmic reticulum

To determine where RG10 acts within the cell, we generated a fluorescent version of this molecule which was coupled to a small, cell permeable, fluorescent Rhodamine dye (Figure 3A). The RG10-rhodamine (RG10-Rhod) compound retained antiviral activity, although higher doses were required (Figure 3B). By confocal microscopy, we found that RG10-Rhod colocalizes with the endoplasmic reticulum (ER) resident protein Calnexin (Figure 3C). In contrast, free Rhodamine clustered away from ER structures (Figure 3C), suggesting that ER distribution of RG10-Rhod is conferred by its RG10 moiety. Using suboptimal RG10-Rhod concentrations allowing low levels of CoV2 replication, we found that both RG10-Rhod and virus replication complexes localize to the ER, although in segregated regions (Figure 3D). This suggests that RG10 does not act directly onto viral RNA replication and is rather a host-targeting agent (HTA).

**Figure 3.**
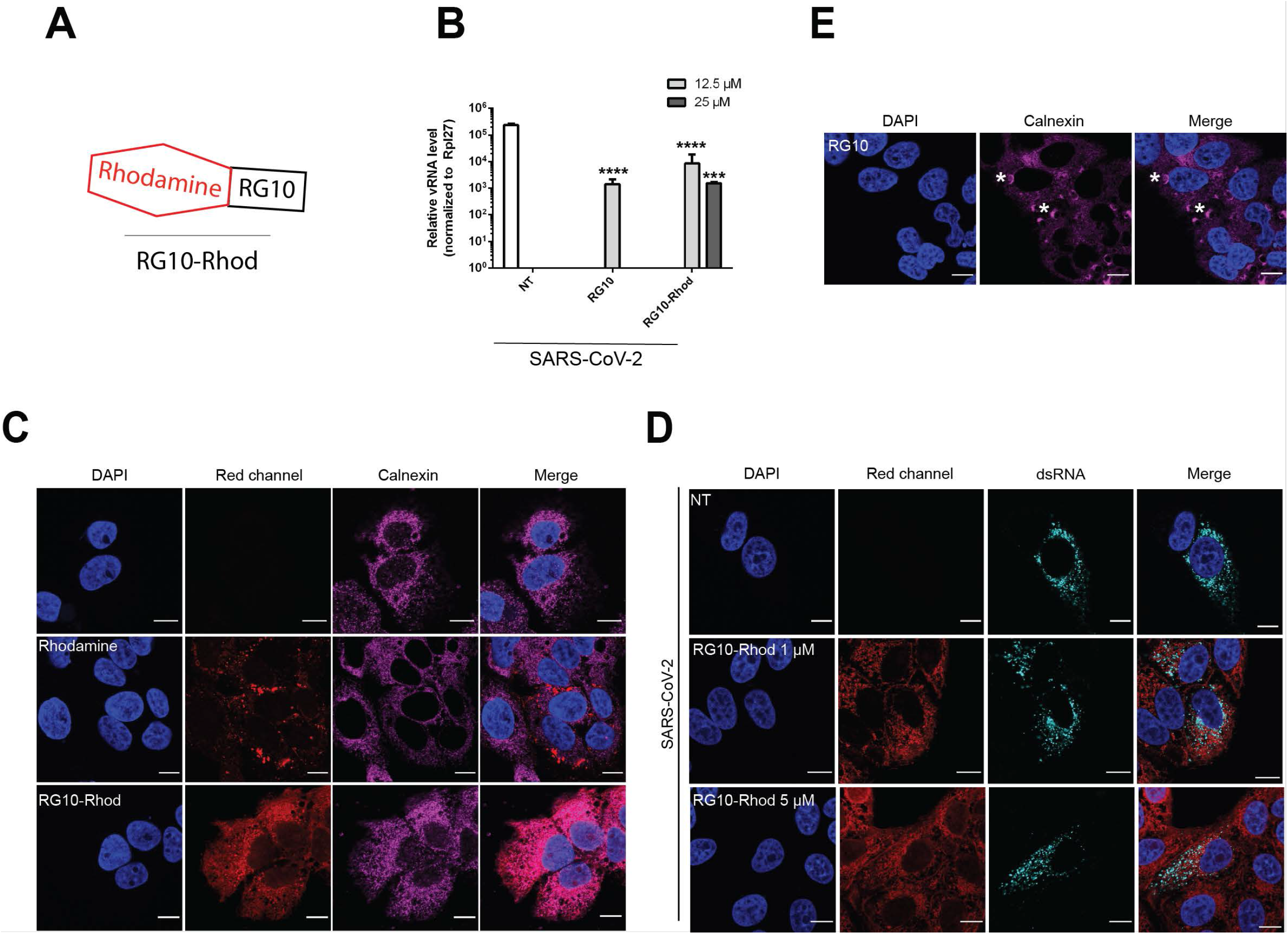
RG10 localize to the endoplasmic reticulum, apart from SARS-CoV-2 replication sites. **(A)** Schematic representation of RG10 molecule coupled to fluorescent Rhodamine (RG10-Rhod) consisting of covalently linked RG10 and Rhodamine moieties. **(B)** Vero E6 cells were treated with indicated concentrations of RG10 or RG10-Rhod and subsequently infected with SARS-CoV-2 at MOI 0.01 for 48 h in the presence of the compounds. The graph shows relative vRNA levels measured by RT-qPCR and normalized by Rpl27 RNA expression. The bars are means +/- SD from duplicates representative of two independent experiments. Student’s t-test *** p value < 0.005; **** p value < 0.001. **(C)** Huh7.5.1 cells were treated with 25 µM of RG10-Rhod or free Rhodamine for 3 h. Cells were fixed and stained using an anti-Calnexin antibody as a marker for ER membranes (magenta) and DAPI for nuclear staining (blue). The red channel corresponds to rhodamine fluorescence. The micrographs show the individual channels and merge that have been acquired using confocal microscopy and processed using ImageJ. Scale bar = 10 µm. **(D)** Vero E6 cells were treated with 1 or 5 µM of RG10-Rhod and then were infected with CoV2 for 24 h in the presence of the inhibitors. Samples were fixed and dsRNA staining was performed to localize viral replication complexes. **(E)** Huh7.5.1 cells were treated with 50 µM of RG10 for 6 h and then were processed for confocal imaging.

### High concentration of RG10 induces sXPB-1-mediated ER stress

We observed that RG10 perturbed ER morphology (Figure 3E and Suppl Figure S1A) and thus wondered whether RG10 could exert its antiviral activity through ER stress induction. Hence, we measured the levels of RNAs of known ER stress mediators from the IRE1-α and PERK pathways and found that the spliced form of XBP-1 (sXBP-1) was enriched upon 50 µM RG10 and 1 mM DTT treatment (Supplemental Figure S1B). In contrast, 50 µM of the RG10 derivative compound RG7, which has no antiviral activity (Figure 1A) did not significantly increase sXBP-1 mRNA levels (Supplemental Figure S1C). However, the increase of sXBP-1 RNA upon RG10 treatment was lower at 10 µM and no difference was observed between RG10 and RG7 (Supplemental Figure S1C), and thus, ER stress induction cannot fully explain the antiviral activity of RG10.

### Generation of infectious CoV2 fluorescent particles

To gain further insights into the mode of action of RG10 antiviral activity, we generated fluorescent infectious CoV2 using a trans-complementation approach. To this end, we generated a Vero E6 cell line stably expressing the nucleocapsid (N) from CoV2 fused to the red fluorescent protein mRuby3. Wild type or Vero E6 N-mRuby3 cells were infected with wild type CoV2 virus (Supplemental Figure S2A-B) and the produced virions incorporated a subset of fluorescent N-mRuby3 proteins in *trans* (not encoded by the viral genome; Figure 4A). The N-mRuby3-containing viruses were as infectious as their non-fluorescent counterparts (Figure 4B and Supplemental Figure S2C), indicating that they represent suitable tools to study CoV2 infection.

**Figure 4.**
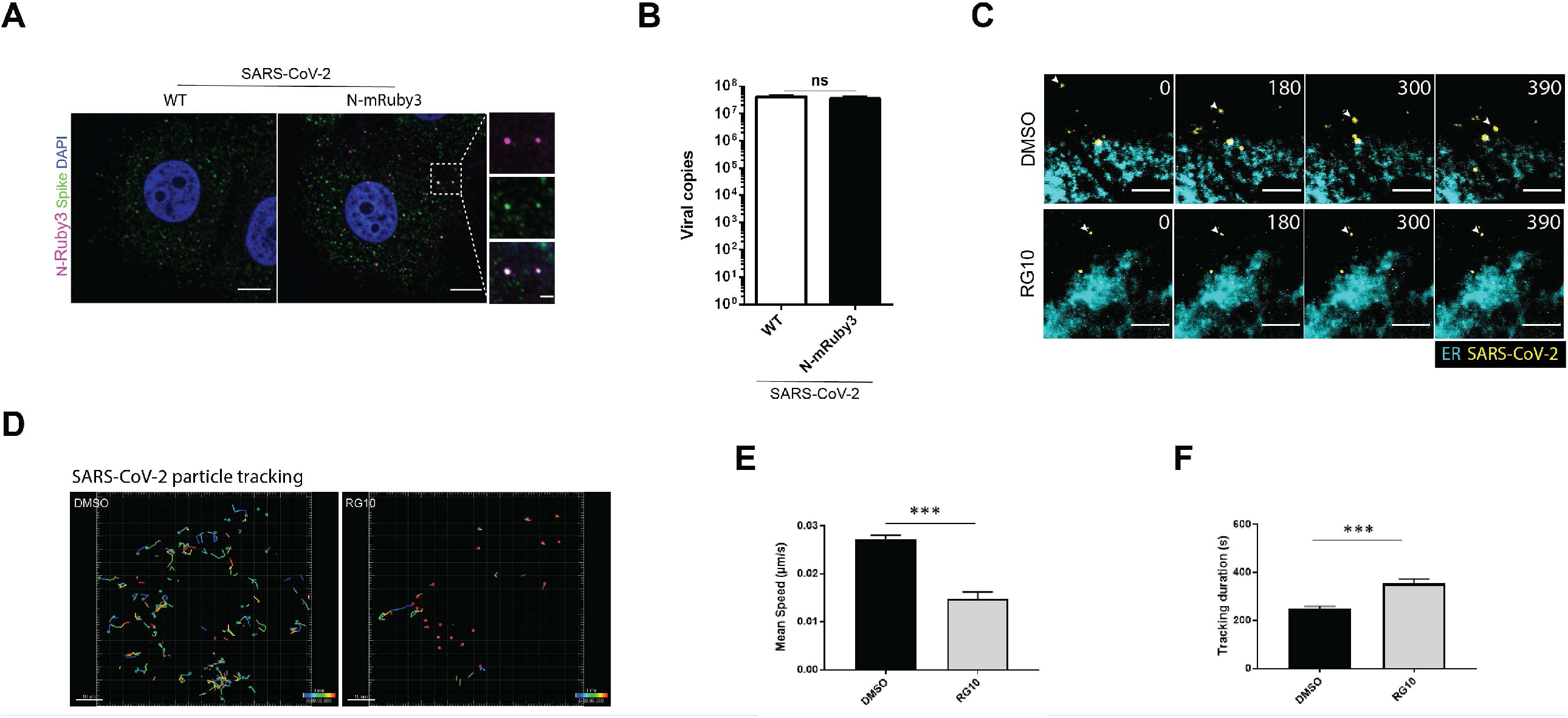
RG10 perturbs virus and ER dynamics. **(A)** Vero E6 cells were infected with WT or N-mRuby3 CoV2 for 3 h at MOI 1. Cells were fixed, permeabilized and stained with DAPI (blue) and an anti-spike antibody (green). Images are from a single z plan acquired by spinning disk confocal microscopy. Scale bar = 10 µm. Magnified images from the dashed square highlight the co-distribution of N-mRuby3-containing virus particles (magenta) and Spike. The zoomed region is represented as single and merge images. Scale bar = 2 µm. **(B)** Vero E6 or Vero E6 N-mRuby3 expressing cells were infected with CoV2 for 3 days. The graph shows relative supernatant vRNA levels measured by RT-qPCR and normalized by Rpl27 RNA expression. **(C)** Huh.7.5.1 cells were labeled for the endoplasmic reticulum using the ER painter reagent and then infected with N-mRuby3 CoV2 at MOI 1. Cells were either incubated with DMSO or 10 µM of RG10 during infection. The micrographs represent snapshots from Supplemental Movies S1. The white arrows highlight a single particle over time for each condition. Scale bar = 10 µm. **(D-F)** CoV2 particles were individually segmented using the same intensity threshold for all conditions and particles were tracked using the Imaris software. **(D)** Representative tracking trails in DMSO or RG10-treated cells associated to Supplemental Movies S2. Particles are in red and each track is color-coded for temporal visualization. The RG10 condition shows viral particles with barely any visible track because the viruses were extremely steady. Quantification of the mean speed **(E)** and tracking duration **(F)** are shown for n = 8 movies per conditions with > 500 virus per condition. Student’s t-test *** p value < 0.005.

### RG10 perturbs ER-CoV2 dynamics

Live cell imaging of CoV2 N-mRuby3 viral particles infecting cells pre-treated with ER painter, a live fluorescent ER marker, allowed us to monitor infection dynamics in untreated and RG10-treated cells (Figure 4C and Movie S1). Spatiotemporal tracking of viral particles showed that the virus was stalled upon RG10 treatment (Figure 4D and Movies S2). Quantification of the mean particle speed confirmed that RG10 slowed down virus movement to the ER (Figure 4E). Because the RG10-treated cells were more immobile, they were more reliably tracked, resulting in higher track duration measurement (Figure 4F). These data indicate that RG10 perturbs intracellular virus dynamics and its transport toward the ER.

### RG10 exhibits coronavirus specific antiviral activity

Next, we aimed to determine whether RG10 had antiviral activity against another virus known to replicate at the ER. Therefore, we compared in parallel the infection of Vero E6 cells by CoV2 or by Zika virus (ZIKV), another enveloped positive-strand RNA virus but from the Flaviviridae family. Interestingly, we found that RG10 had no detectable effect on ZIKV, while in contrast, the ER stress inducer Tunicamycin was active against both viruses (Figure 5A-B). To determine whether RG10 was specific to CoV2 or can act against other coronaviruses, we tested its activity against the human coronavirus 229E (HCoV-229E) and observed significant antiviral activity, although potency was lower (Figure 5C).

**Figure 5.**
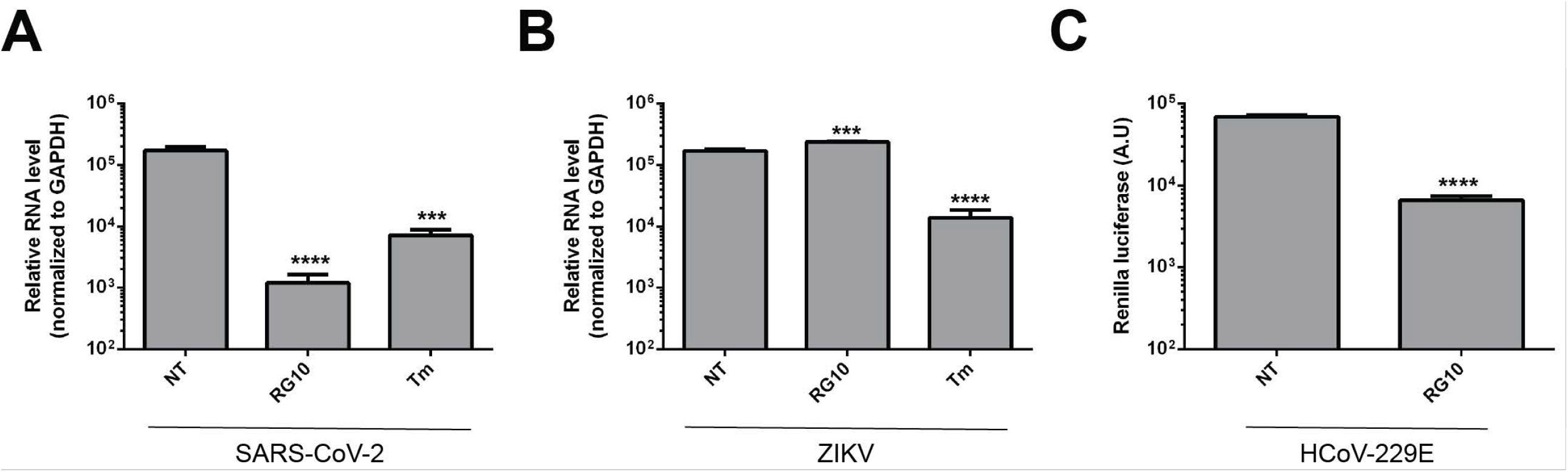
RG10 exhibits pan-coronavirus activity but does not inhibit Zika virus infection. **(A-B)** Vero E6 cells were treated with 10 µM RG10 or 1 µM Tunicamycin (Tm) and infected with CoV2 at MOI 0.01 (A) or ZIKV at MOI 0.3 (B) for 24 h in the presence of the compounds. The graph shows relative vRNA levels measured by RT-qPCR and normalized by GAPDH RNA expression. The bars are means +/- SD from duplicates representative of two independent experiments. Student’s t-test *** p value < 0.005; **** p value < 0.001. **(C)** Huh7.5.1 cells were treated with 10 µM RG10 and infected with HCoV-229E-Renilla at MOI 2 for 24 h in the presence of the compound.

### RG10-mediated inhibition of CoV2 in human airway epithelia

To move toward effective preclinical assays, we tested the efficacy of RG10 in human airway epithelia (HAE), an *ex vivo* model of primary cells cultured in 2D on transwells, exhibiting expected cilia beating activity. In this model, we found that RG10 decreases viral infection using a fluorescent mNeonGreen replication-competent virus (Xie et al., 2020), and that virus protein expression was also altered (Figure 6A-B). Although HAE viability was similar, the actin network seemed to be rearranged at the apical side of RG10-treated HAE (Figure 6C). Interestingly, sXBP-1 was upregulated upon RG10 treatment (Figure 6D), indicating that RG10 mediates ER stress in this primary HAE model. By RT-qPCR, viral RNA was decreased by RG10, although the extent of the decrease was much lower than Remdesivir treatment (Figure 6E).

**Figure 6.**
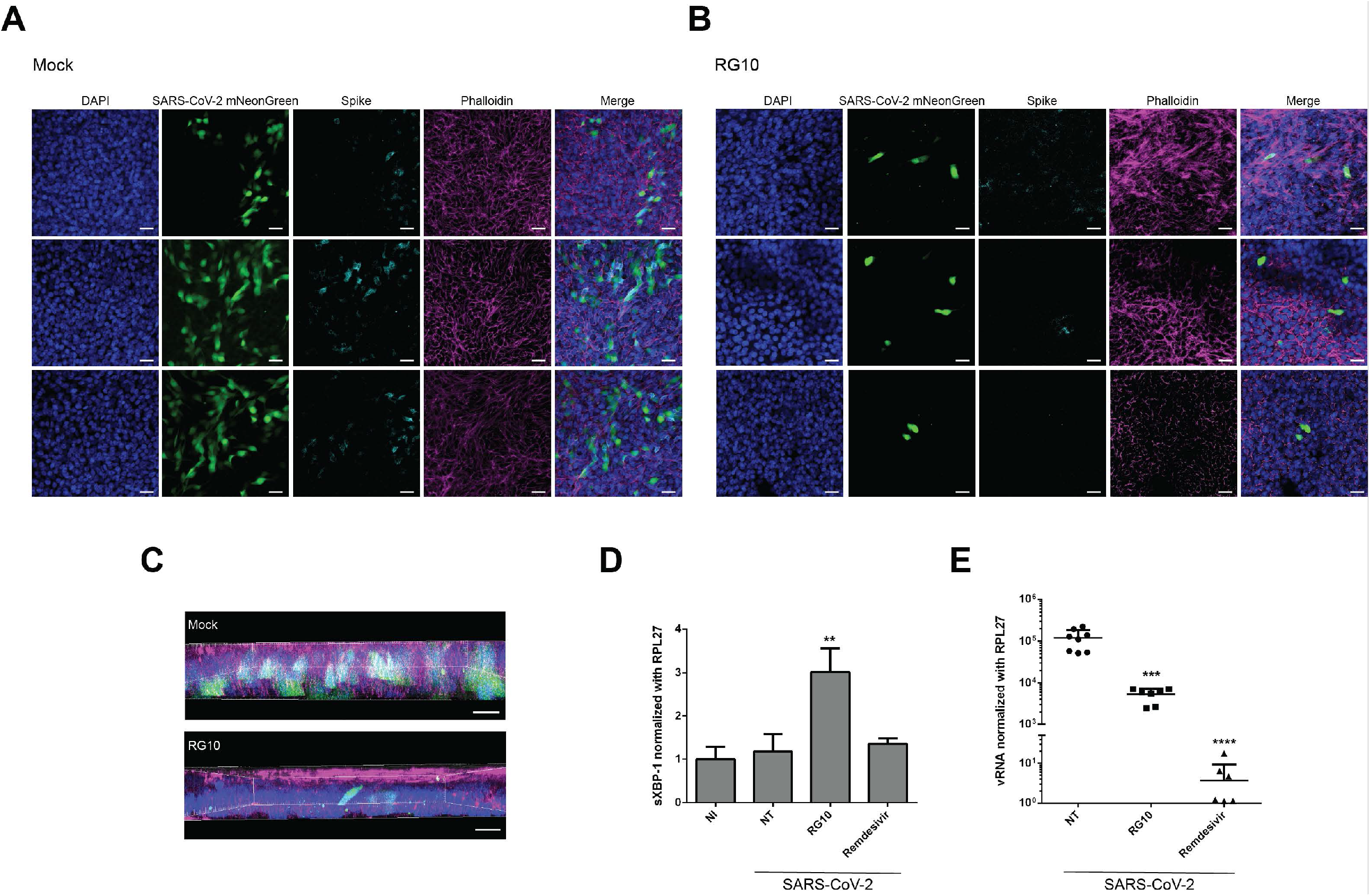
RG10 prevents SARS-CoV-2 infection of human airway epithelia. **(A-B)** HAE cells were either untreated (Mock) or treated with 10 µM of RG10 and then infected with SARS-CoV-2 mNeonGreen virus at MOI 0.1. After 4 days, cells were fixed and stained for CoV2 spike protein and actin filaments (phalloidin staining). Z-stack images were acquired using a confocal microscope and processed using ImageJ to produce an average intensity image for each individual channel. Scale bar = 20 µm. **(C)** 3D representation of the images from **A-B** using the Imaris software. Scale bar = 20 µm. **(D)** HAE cells were treated with 10 µM of RG10 and Remdesivir and then infected with wild type CoV2 at MOI 0.01 for 4 days. Graphs show relative sXBP1 RNA levels normalized with GAPDH RNA expression. **(E)** Same samples from **D** were processed to measure by RT-qPCR, relative vRNA levels normalized by Rpl27 RNA expression. The bars are means +/- SD from two pooled independent experiments. Student’s t-test ** p value < 0.01; *** p value < 0.005; **** p value < 0.001.

### Identification of a more potent and stable RG10 derivative

The low potency of RG10 in HAE could be attributed to its weak stability and therefore, we designed a novel RG10 derivative, which we called RG10b. This derivative harbors modifications shielding a molecular weakness present in RG10. RG10b was more potent than RG10 with an EC_50_ < 1 µM (Figure 7A-B). To verify the stability of RG10 and RG10b in physiological conditions, we performed pharmacokinetics analyses in mice and found that RG10b has a half-life of 2h19 in the blood, a value more than 3-fold higher than for RG10 (Figure 7C). Innocuity assessment in mice showed that RG10b was well supported by mice at concentrations up to 10 mg/kg (higher concentrations were not tested; Figure 7D). Treatment of HAE cells with 10 µM of the indicated compounds confirmed that RG10b was more potent than RG10, showing antiviral activity similar to Remdesivir in this *ex vivo* model (Figure 7E).

**Figure 7.**
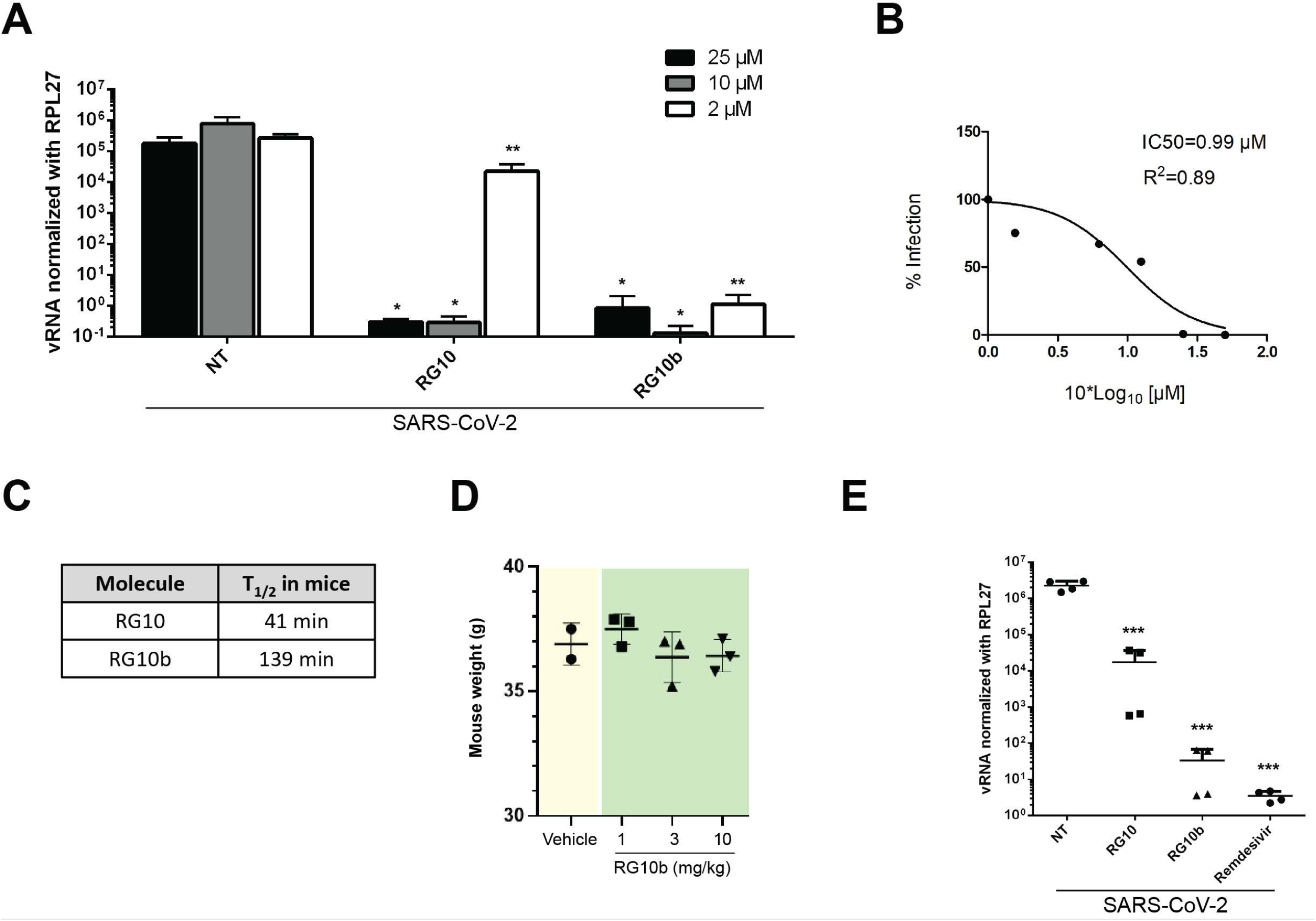
RG10b shows higher potency and stability than RG10 against CoV. **(A)** Vero E6 cells were treated with the indicated concentrations of RG10 or RG10b and then infected with CoV2 for 48 h. Graphs show relative vRNA levels normalized with Rpl27 RNA expression. **(B)** Vero E6 cells were treated with RG10b at different concentrations and subsequently infected and processed as in **A**. The IC_50_ of RG10b was ≈ 0.99 µM. **(C)** Table shows T_1/2_ of RG10 and RG10b as determined by pharmacokinetics experiments in mice. **(D)** Graph shows weight of mice after treatment with various concentrations of RG10b. Two mice were used per condition. **(E)** HAE cells were treated with 10 µM of RG10, RG10b, and Remdesivir and then infected for 4 days with CoV2 at MOI 0.01. Cells were then processed for RNA extraction and RT-qPCR experiments. Graph shows relative vRNA levels normalized with Rpl27 RNA expression. Student’s t-test * p value < 0.05 ** p value < 0.01; *** p value < 0.005.

## Discussion

In this work, we have characterized a novel SARS-CoV-2 antiviral molecule and explored its mode of action. To this end, we generated novel tools to study the spatiotemporal dynamics of the virus at high resolution.

Numerous antiviral drugs have been considered for repurposing to treat COVID-19 (for review, see (Indari et al., 2021)). Direct-acting antiviral agents (DAA) targeting the virus usually exhibit very good EC_50_ relative to their toxicity, but RNA viruses being prone to mutations (as highlighted by the emerging variants) suggest that this approach may not be sufficient by itself to counteract virus dissemination and may instead favor escape mutants in monotherapies. In contrast, HTAs that target cellular partners important for the virus life cycle is an approach of choice, that must be considered in parallel to DAA and vaccination. Here, we chose a bottom-up approach, using a series of related proprietary molecules that had no known functions, and started to characterize it. During this study, we also generated new tools to perform live cell imaging of SARS-CoV-2 entry, offering the opportunity to learn more about the virus’ life cycle, while tailoring an antiviral molecule with promising pharmacological properties.

Approved HTAs often have broad-spectrum antiviral activity due to the fact that viruses may exploit similar pathways. For instance, (hydroxy)chloroquine, ribavirin, and favipiravir are examples of broad-spectrum antivirals that are used *in vitro* at high concentrations to be effective (Driouich et al., 2021, Maisonnasse et al., 2020, Yao et al., 2020, Zhao et al., 2021). Although favipiravir shows very low potency in vitro (EC_50_ > 200 µM in Vero E6 cells), it can decrease CoV2 viral loads in hamsters (Driouich et al., 2021). Hydroxychloroquine, which has an EC_50_ ≈ 2 µM in Vero E6 cells, does not decrease the viral loads of infected non-human primates (Maisonnasse et al., 2020). The heterogeneity between *in vitro* and *in vivo* potency highlights the complexity to develop HTAs. RG10 is likely an HTA as it does not localize at virus replication sites to exert its antiviral activity. The improved version of RG10, RG10b, shows EC_50_ below 1 µM, high stability *in vivo* (> 2 h), with no toxicity observed (Figure 7). These data are promising and future work will aim at assessing the antiviral activity of RG10 and derivatives in animal models.

At this stage, the target of RG10 is unknown but we identified that the molecule strongly affects virus and ER dynamics (Figure 4), suggesting that it perturbs players involved in intracellular trafficking routes. Concordantly, we showed that the actin cytoskeleton was disorganized by RG10 in primary HAE cells (Figure 6). Preserved integrity of the actin cytoskeleton is required for SARS-CoV-2 infection (Yeung et al., 2021, Zhang et al., 2020). The FDA-approved drugs Sunitinib and BNTX inhibit SARS-CoV-2 entry and their mode of action was correlated to the disruption of actin dynamics (Zhang et al., 2020). Moreover, SARS-CoV-2 increases filopodia number and length, while traveling along them, further indicating that a strong interplay exists between CoV2 and the actin network (Bouhaddou et al., 2020). RG10 has very low cytotoxicity compared to Latrunculin, jasplakinolide and cytocalasin D, three well-known direct-acting actin perturbators that inhibit CoV2 infection (Yeung et al., 2021, Zhang et al., 2020, Zhang et al., 2021). It is therefore likely that RG10 subtly impacts the actin cytoskeleton in a direct or indirect manner that remains to be identified.

RG10 induces ER stress at high doses, but it was difficult to correlate this observation to the antiviral effect of RG10 because at lower doses, RG10 and RG7, two closely related molecules, similarly induced ER stress (Supplemental Figure S1) even though RG7 showed no antiviral activity (Figure 1). Furthermore, RG10 was unable to inhibit ZIKV replication, which also occurs in the ER. We used Tunicamycin as a potent ER stress inducer that inhibits infection by CoV2 and ZIKV (our data and (Mufrrih et al., 2021)), further suggesting that ER stress induction may not be sufficient for RG10 to exert an antiviral activity.

We found that RG10 is targeted to the ER and impacts ER morphology. Constant reshaping of the ER structure is a requirement for maintaining its numerous functions, and ER membranes are often targeted by pathogens (Ravindran et al., 2016). For instance, influenza A virus was recently shown to impact mitochondrial morphodynamics and endoplasmic reticulum-mitochondria contact sites, and altering mitochondrial dynamics was thus proposed as a new antiviral strategy (Pila-Castellanos et al., 2021). Similarly, one can hypothesize that ER morphodynamics can represent an antiviral strategy to be further explored. In this regard, atlastins play important roles in ER dynamics and morphology and it was recently shown that atlastin-dependent misshaped ER leads to decreased ZIKV replication (Monel et al., 2019). To our knowledge however, the importance of atlastins during coronavirus infection has not been investigated so far. In any case, ER morphology seems to be important for CoV2 infection, and perturbation of ER dynamics by RG10 is likely to strongly contribute to its antiviral activity.

The study of CoV2 spatiotemporal dynamics, including virus entry, replication, assembly, and release requires new tools to quantitatively assess it. A comprehensive list of plasmids coding for the CoV2 viral proteins fused to fluorescent proteins has been generated, which proved very useful to evaluate the distribution and potential function of individual viral proteins (Miserey-Lenkei et al., 2021). Moreover, fluorescent virus-like particles (VLP) expressing Spike from CoV2 were generated, although with modest specificity (Ma et al., 2021), but to date, fluorescent viral particles containing the full wild type genome of CoV2 has not been described. Using a trans-complementation approach, we generated bright CoV2 particles which retain infectiveness comparable to their wildtype counterparts. Although we exploited this reagent to study CoV2 arrival to ER membranes, fluorescent viral particles shall prove useful in a variety of assays aiming at a better understanding of CoV2 entry.

## Material & Methods

### Antibodies and reagents

Mouse anti-Spike and anti-GAPDH were purchased from Genetex and Rabbit anti-N from Sino Biological (Clinisciences). The mouse anti-dsRNA antibody (J2) was from Jena Bioscience and rabbit anti-Calnexin from Elabscience. NHS-Rhodamine and all secondary Alexa Fluor antibodies were purchased from Thermo Fisher Scientific. Phalloidin CF-633 was purchased from Biotium and the ER Staining Kit was from Abcam. RG10 and derivatives were synthesized by AGV Discovery.

### Cell lines and primary cells

Huh7.5.1 (Zhong et al., 2005), Vero E6 (ECACC #85020206), Caco2, and HEK 293T cells were cultured in DMEM GlutaMAX (Thermo Fisher Scientific) supplemented with 10% fetal calf serum and penicillin-streptomycin (500 μg/ml; GIBCO). HAE cells (Epithelix) were cultured in Mucilair media at the basal side and maintained at air-liquid interface. Apical washes were performed each 2-3 days by temporarily adding Mucilair media for 30 minutes. All cell types were maintained at 37 °C in a 5% CO2 atmosphere.

### Plasmids

To generate a lentiviral vector containing the Nucleocapsid (N) of SARS-CoV-2 fused in C-terminus to the gene coding for the monomeric red fluorescent protein mRuby3 (Bryce T. Bajar et al, Sci Rep, 2016), N was firstly amplified by PCR from the plasmid pLVX-EF1alpha-SARS-CoV-2-N-2xStrep-IRES-Puro (addgene#141391) using the sense primer 5’ TTGC<u>GGCCGC</u>GCCACCATGAGCGATAACG-3’ (NotI site underlined) and the antisense primer 5’-CGCCTGAGTAGAATCGGCT-3 (overlapped region underlined). The Ruby-3 fragment was amplified by PCR from a gblock (Integrated DNA-Technologies) using the sense primer 5’-AGCCGATTCTACTCAGGCGGGAGGTTCTGGTGGTTCTG-3’ (overlapped region underlined) and the antisense primer 5’-GA<u>GGATC</u>CTCACTTGTACAGCTCGTCCAT-3’ (BamHI site underlined). The individual N and Ruby3 fragments were linked by overlapping PCR of the N-CoV2 using primers 5’-TTGCGGCCGCGCCAC-3’ and 5’-GAGGATCCTCACTTGTACAGCTCGT-3’. The resulting PCR product was ligated into the digested NotI and BamHI lentiviral plasmid pHAGE IRES puro. The resulting plasmid pHAGE N-mRuby3 IRES puro is available on Addgene (#170466). In parallel, a pHAGE N-mNeonGreen IRES puro was also generated (Addgene #170467).

### Generation of stable cell line

HEK 293T cells were transfected with pxPAX2, VSV-G and pHAGE N-mRuby3 IRES puro using JetPrime (Poluplus Transfection) and washed 6 h post-transfection. The supernatant was harvested 2 days later and cleared by centrifugation at 2000 x g for 5 min at 4°C. The lentivirus-containing supernatant was added to Vero E6 cells (ECACC #85020206) for 48 h. Selection of the positive cells was performed using 4 µg/ml puromycin and viable cells were expanded for flow cytometry cell sorting using a FACSMelody (BD Biosciences) equipped with a 561 nm excitation laser and 613/18 filter. Upon sorting, the cells were tested negative for mycoplasma. Vials of the Vero E6 N-Ruby3 stable cell line were conserved in liquid nitrogen in 10% DMSO.

### Virus production and titration

The SARS-CoV-2 strain BetaCoV/France/IDF0372/2020 was supplied by the National Reference Centre for Respiratory Viruses hosted by Institut Pasteur (Paris, France) and headed by Dr. Sylvie van der Werf. The strain was amplified through the infection of Vero E6 cells at MOI 0.0001 in DMEM GlutaMAX (Gibco) supplemented with 2% fetal bovine serum (FBS; Dominique Dutscher) and 1X penicillin-streptomycin (GIBCO). At 3 days post infection (dpi), the supernatant was harvested and cleared by centrifugation at 2000 x g for 5 min at 4°C. The cleared virus-containing supernatant was frozen in 1 ml aliquots at -80°C. For each virus production, a vial was thawed for titration by plaque assay in Vero E6 cells to estimate plaque-forming units per mL of virus (PFU/mL) as described in Gaudin & Barteneva (Gaudin & Barteneva, 2015). Viral titers ranged between 3 10^6^ and 2 10^7^ pfu/ml.

HCoV-229E-Renilla was a gift from Volker Thiel (van den Worm et al., 2012) and was amplified for 5-7 days at 33°C in Huh7.5.1 cells in 5% FCS-containing DMEM. Viral stocks were harvested when cells showed > 50% CPEs. Virus stock was titrated through TCID_50_ in Huh7.5.1 cells used for their amplification and titer was 10^8^ TCID_50_/mL. Infections of 50 000 cells were performed at MOI 2 for HCoV-229E-Renilla (as measured on Huh7.5.1 cells) and infection efficiency was analyzed one day later by measuring Renilla activity (Renilla Luciferase Assay System, Promega).

The ZIKV strain was obtained through BEI Resources (NIAID NR-50183, strain FLR) isolated in December 2015 from infected patient blood from Colombia. ZIKV was amplified in Vero cells for 72 h in DMEM GlutaMAX (Gibco) supplemented with 2% fetal bovine serum (FBS; Dominique Dutscher), 10 mM HEPES (Thermo Fisher Scientific) and 1X penicillin-streptomycin (Gibco). The supernatant was harvested and clarified by centrifugation at 2000 g RT for 10 min. The virus was stored at -80°C. The virus titer was obtained by using standard plaque assay on Vero cells to measure plaque-forming unit per mL of virus (PFU/mL). Viral titer was 3.8 ×10^6^ pfu/mL.

### RT-qPCR

Cells were lysed and RNA immediately extracted using the Luna Cell Ready Lysis Module according to the manufacturer’s protocol (New England Biolabs). RT-qPCR on extracted RNA was performed using the Luna One-Step RT-qPCR Kit (New England Biolabs). The samples were analyzed using a Lightcycler 480 instrument (Roche) using the following thermal cycle: Reverse transcription step at 55 °C for 15 min, initial denaturation at 95 °C for 2 min. Then, 40 cycles were programmed consisting of 15 s of denaturation at 95 °C followed by 30 s of annealing/extension at 55 °. Fluorescent data collection was performed at the end of each cycle. SARS-CoV-2 viral RNA was detected using the N1 probe that targets specifically the N protein. For normalization, primers targeting Rpl27 or GAPDH were utilized.

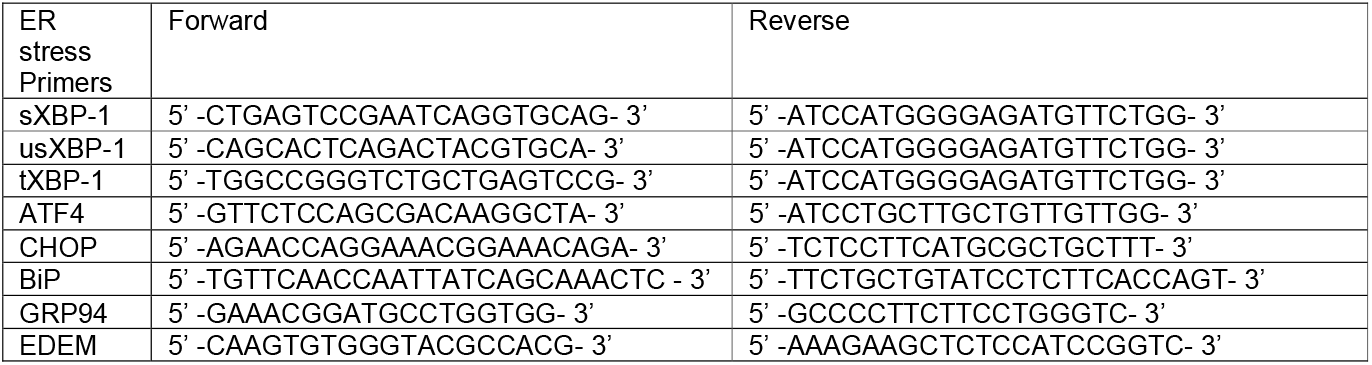

### Immunofluorescence

Cells grown on #1.5 glass coverslips were fixed with 4 % PFA for 15 min. Cells were then permeabilized with 0.1% Triton ×100 and saturated with 0.2% BSA. Then, primary antibody incubation was performed for 1 h. After a series of PBS washes, secondary antibodies were added for 30 min. DAPI staining was performed for 10 min to obtain nuclear staining. Coverslips were mounted on glass slides using Mowiol mounting media. Images were acquired using a Leica SP5 confocal microscope.

### Live cell imaging

Image acquisition was performed on an AxioObserver.Z1 inverted microscope (Zeiss) mounted with a CSU-X1 spinning disk head (Yokogawa), a back-illuminated EMCCD camera (Evolve, Photometrics) and a ×100, 1.4 NA oil objective (Zeiss) controlled by MetaMoprh. The microscope chamber was heated at 37°C and the automated multiposition stage maintained in 95% humidity at 37°C and 5% CO_2_. Images were processed using Fiji (ImageJ software version 1.51h) and quantitative analyses were performed using Bitplane Imaris x64 version 9.2. Analyses of the movies consisted of the segmentation of the single viral particles based on intensity and size. Then, tracking was performed for each individual particle and mean speed and track duration were extracted.

### Statistical analysis

Statistical analyses of the data were performed using two-tailed unequal variance Student t-tests (ns p>0.05, * p<0.05, ** p<0.01, *** p<0.001) unless mentioned otherwise. The mean and standard deviation (SD) of the mean were plotted using GraphPad Prism version 7.04. Number of independent experiments (n) is indicated in the figure legends.

## Acknowledgments

We thank Pr. Sylvie van der Werf and the National Reference Centre for Respiratory Viruses hosted by Institut Pasteur (Paris, France) for providing us with the wild type SARS-CoV-2 strain and Dr. Pei-Yong Shi (Galveston, TX, USA) for the SARS-CoV-2 mNeonGreen reporter virus (Xie et al., 2020). We acknowledge the imaging facility MRI, member of the national infrastructure France-BioImaging and the region BSL-3 research facilities CEMIPAI for providing excellent infrastructures to work on infectious SARS-CoV-2. This work was funded by CNRS-INSB and SATT-AxlR to R.G. We would like to acknowledge the Drug discovery center (PCBIS, Illkirch, France) for in vivo PK and innocuity analyses.

## Supplemental information

**Supplemental Figure S1.**
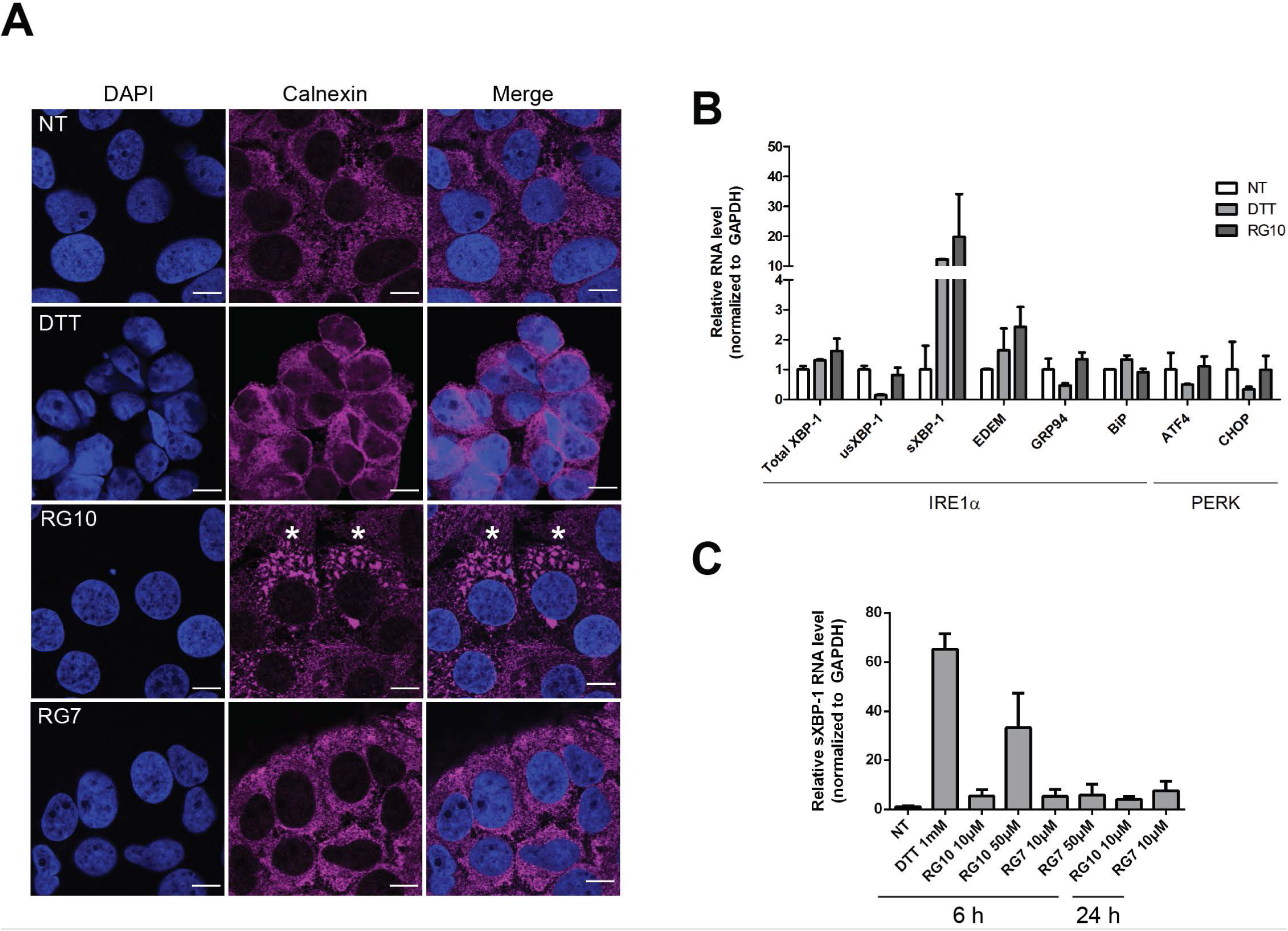
RG10 induces ER stress in a sXBP-1-dependent manner. **(A)** Huh7.5.1 cells were treated with 50 µM RG7 (an inactive derivative of RG10), 50 µM RG10 or 1 mM DTT for 6 h. Cells were then fixed and stained using an anti-Calnexin antibody as a marker for ER membranes (magenta) and DAPI for nuclear staining (blue). The micrographs show the individual channels and merge that have been acquired using confocal microscopy and processed using ImageJ. Scale bar = 10 µm. **(B)** Huh7.5.1 cells were treated as in A. The graph shows relative RNA levels measured by RT-qPCR and normalized by GAPDH RNA expression. The bars are means +/- SD from duplicates representative of two independent experiments. **(C)** Huh.7.5.1 cells were treated with either 10 µM or 50 µM of RG10 or RG7 for 6 or 24 h. Exposure for 6 h to 1 mM of DTT was used as a positive control of ER stress. The graph shows relative spliced sXBP-1 levels measured by RT-qPCR and normalized by GAPDH RNA expression.

**Supplemental Figure S2.**
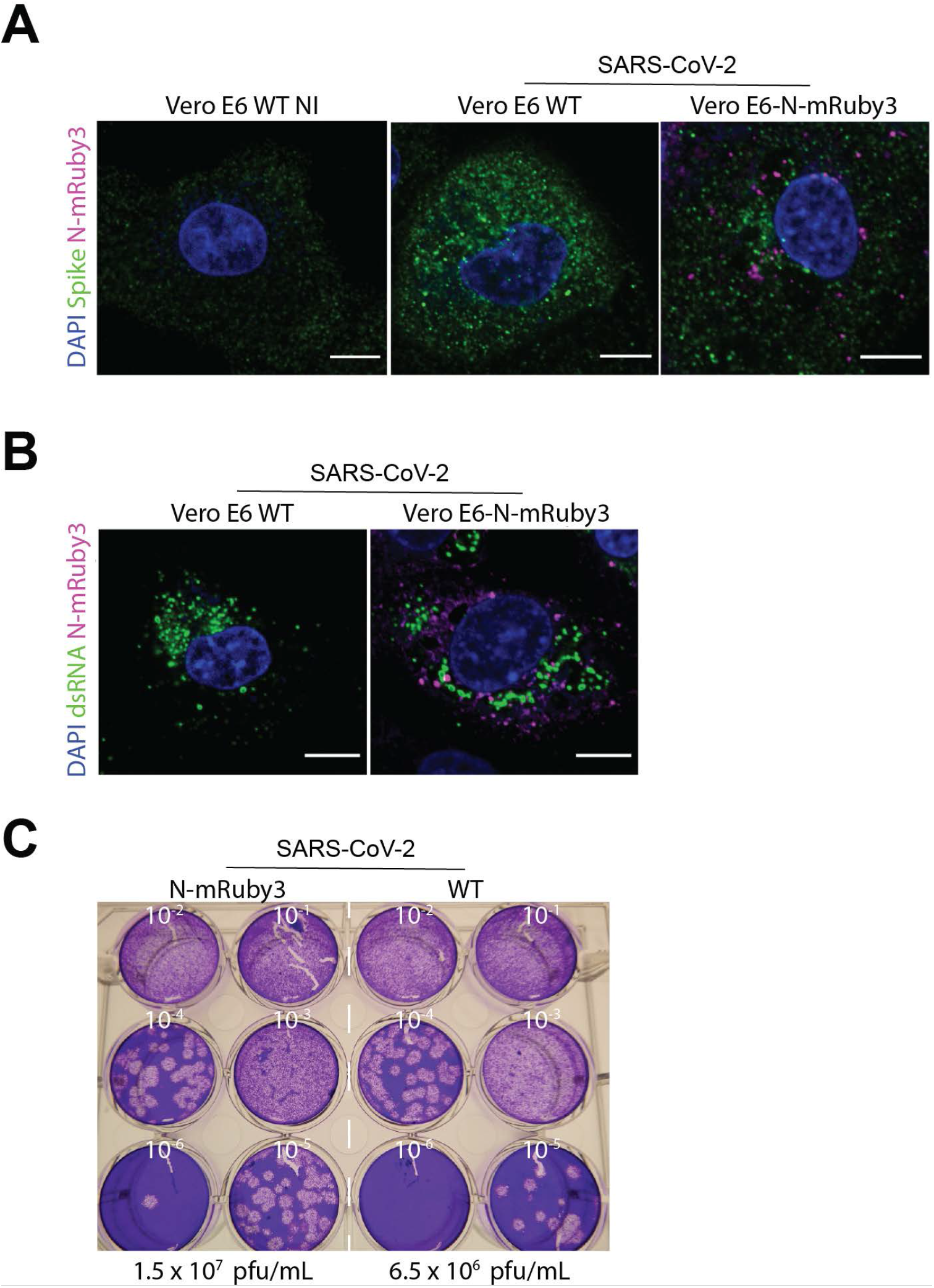
Characterization of fluorescent SARS-CoV-2 N-mRuby3 viral particles. **(A-B)** Vero E6 WT cells and Vero E6 cells stably expressing N-mRuby3 were infected for 48 h at MOI 0.01 with CoV2. Cells were fixed, permeabilized and stained with DAPI (blue) and either an anti-Spike antibody (green, **A**) or an anti-dsRNA antibody (green, **B**). N-mRuby3 is shown in magenta. Merged immunofluorescence images are from a single z plan acquired by spinning disk confocal microscopy. Scale bar = 10 µm. **(C)** Vero E6 or Vero E6 N-mRuby3 expressing cells were infected with CoV2 for 3 days. Supernatants were then used to perform plaque assays on Vero E6 cells.

**Supplemental Movie S1. Effect of RG10 on intracellular trafficking of fluorescent SARS-CoV-2 N-mRuby3 viral particles**. Huh7.5.1 cells were labeled with ER painter (cyan) and treated with DMSO (Mock, left) or RG10 (right) and then infected with fluorescent SARS-CoV-2 N-mRuby3 viral particles (yellow) at MOI 1 for 2 h. Cells were imaged every 30 s for 10 min using spinning disk confocal microscopy.

**Supplemental Movie S2. Single particle tracking of SARS-CoV-2 in the presence or absence of RG10**. The movies show the tracking of SARS-CoV-2 N-mRuby3 particles (red) under DMSO-treated (Mock, left) or RG10-treated (right) conditions. The color-coding of the tracks indicates the position of the particles over time. Cells were imaged every 30 s for 10 min using spinning disk confocal microscopy.

## References

Bayati A, Kumar R, Francis V, McPherson PS (2021) SARS-CoV-2 infects cells following viral entry via clathrin-mediated endocytosis. J Biol Chem: 100306

Bouhaddou M, Memon D, Meyer B, White KM, Rezelj VV, Correa Marrero M, Polacco BJ, Melnyk JE, Ulferts S, Kaake RM, Batra J, Richards AL, Stevenson E, Gordon DE, Rojc A, Obernier K, Fabius JM, Soucheray M, Miorin L, Moreno E et al. (2020) The Global Phosphorylation Landscape of SARS-CoV-2 Infection. Cell 182: 685–712 e19

Driouich JS, Cochin M, Lingas G, Moureau G, Touret F, Petit PR, Piorkowski G, Barthelemy K, Laprie C, Coutard B, Guedj J, de Lamballerie X, Solas C, Nougairede A (2021) Favipiravir antiviral efficacy against SARS-CoV-2 in a hamster model. Nature communications 12: 1735

Gaudin R, Barteneva NS (2015) Sorting of small infectious virus particles by flow virometry reveals distinct infectivity profiles. Nature communications 6: 6022

Goldman JD, Lye DCB, Hui DS, Marks KM, Bruno R, Montejano R, Spinner CD, Galli M, Ahn MY, Nahass RG, Chen YS, SenGupta D, Hyland RH, Osinusi AO, Cao H, Blair C, Wei X, Gaggar A, Brainard DM, Towner WJ et al. (2020) Remdesivir for 5 or 10 Days in Patients with Severe Covid-19. The New England journal of medicine

Hoffmann M, Kleine-Weber H, Schroeder S, Kruger N, Herrler T, Erichsen S, Schiergens TS, Herrler G, Wu NH, Nitsche A, Muller MA, Drosten C, Pohlmann S (2020) SARS-CoV-2 Cell Entry Depends on ACE2 and TMPRSS2 and Is Blocked by a Clinically Proven Protease Inhibitor. Cell

Huang C, Wang Y, Li X, Ren L, Zhao J, Hu Y, Zhang L, Fan G, Xu J, Gu X, Cheng Z, Yu T, Xia J, Wei Y, Wu W, Xie X, Yin W, Li H, Liu M, Xiao Y et al. (2020) Clinical features of patients infected with 2019 novel coronavirus in Wuhan, China. Lancet 395: 497–506

Indari O, Jakhmola S, Manivannan E, Jha HC (2021) An Update on Antiviral Therapy Against SARS-CoV-2: How Far Have We Come? Front Pharmacol 12: 632677

Ma Y, Mao G, Wu G, Chen M, Qin F, Zheng L, Zhang XE (2021) Dual-Fluorescence Labeling Pseudovirus for Real-Time Imaging of Single SARS-CoV-2 Entry in Respiratory Epithelial Cells. ACS Appl Mater Interfaces 13: 24477–24486

Maisonnasse P, Guedj J, Contreras V, Behillil S, Solas C, Marlin R, Naninck T, Pizzorno A, Lemaitre J, Goncalves A, Kahlaoui N, Terrier O, Fang RHT, Enouf V, Dereuddre-Bosquet N, Brisebarre A, Touret F, Chapon C, Hoen B, Lina B et al. (2020) Hydroxychloroquine use against SARS-CoV-2 infection in non-human primates. Nature 585: 584–587

Miserey-Lenkei S, Trajkovic K, D’Ambrosio JM, Patel AJ, Copic A, Mathur P, Schauer K, Goud B, Albanese V, Gautier R, Subra M, Kovacs D, Barelli H, Antonny B (2021) A comprehensive library of fluorescent constructs of SARS-CoV-2 proteins and their initial characterisation in different cell types. Biol Cell

Monel B, Rajah MM, Hafirassou ML, Sid Ahmed S, Burlaud-Gaillard J, Zhu PP, Nevers Q, Buchrieser J, Porrot F, Meunier C, Amraoui S, Chazal M, Salles A, Jouvenet N, Roingeard P, Blackstone C, Amara A, Schwartz O (2019) Atlastin Endoplasmic Reticulum-Shaping Proteins Facilitate Zika Virus Replication. Journal of virology 93

Mufrrih M, Chen B, Chan SW (2021) Zika Virus Induces an Atypical Tripartite Unfolded Protein Response with Sustained Sensor and Transient Effector Activation and a Blunted BiP Response. mSphere: e0036121

Ou T, Mou H, Zhang L, Ojha A, Choe H, Farzan M (2021) Hydroxychloroquine-mediated inhibition of SARS-CoV-2 entry is attenuated by TMPRSS2. PLoS pathogens 17: e1009212

Pila-Castellanos I, Molino D, McKellar J, Lines L, Da Graca J, Tauziet M, Chanteloup L, Mikaelian I, Meyniel-Schicklin L, Codogno P, Vonderscher J, Delevoye C, Moncorge O, Meldrum E, Goujon C, Morel E, de Chassey B (2021) Mitochondrial morphodynamics alteration induced by influenza virus infection as a new antiviral strategy. PLoS pathogens 17: e1009340

Ravindran MS, Bagchi P, Cunningham CN, Tsai B (2016) Opportunistic intruders: how viruses orchestrate ER functions to infect cells. Nat Rev Microbiol 14: 407–420

V’Kovski P, Kratzel A, Steiner S, Stalder H, Thiel V (2021) Coronavirus biology and replication: implications for SARS-CoV-2. Nat Rev Microbiol 19: 155–170

van den Worm SH, Eriksson KK, Zevenhoven JC, Weber F, Zust R, Kuri T, Dijkman R, Chang G, Siddell SG, Snijder EJ, Thiel V, Davidson AD (2012) Reverse genetics of SARS-related coronavirus using vaccinia virus-based recombination. PloS one 7: e32857

Walls AC, Park YJ, Tortorici MA, Wall A, McGuire AT, Veesler D (2020) Structure, Function, and Antigenicity of the SARS-CoV-2 Spike Glycoprotein. Cell

Wolff G, Melia CE, Snijder EJ, Barcena M (2020) Double-Membrane Vesicles as Platforms for Viral Replication. Trends Microbiol S0966-842X(20)30135-9.

Xie X, Muruato A, Lokugamage KG, Narayanan K, Zhang X, Zou J, Liu J, Schindewolf C, Bopp NE, Aguilar PV, Plante KS, Weaver SC, Makino S, LeDuc JW, Menachery VD, Shi PY (2020) An Infectious cDNA Clone of SARS-CoV-2. Cell host & microbe 27: 841–848 e3

Yao X, Ye F, Zhang M, Cui C, Huang B, Niu P, Liu X, Zhao L, Dong E, Song C, Zhan S, Lu R, Li H, Tan W, Liu D (2020) In Vitro Antiviral Activity and Projection of Optimized Dosing Design of Hydroxychloroquine for the Treatment of Severe Acute Respiratory Syndrome Coronavirus 2 (SARS-CoV-2). Clinical infectious diseases : an official publication of the Infectious Diseases Society of America

Yeung ML, Teng JLL, Jia L, Zhang C, Huang C, Cai JP, Zhou R, Chan KH, Zhao H, Zhu L, Siu KL, Fung SY, Yung S, Chan TM, To KK, Chan JF, Cai Z, Lau SKP, Chen Z, Jin DY et al. (2021) Soluble ACE2-mediated cell entry of SARS-CoV-2 via interaction with proteins related to the renin-angiotensin system. Cell 184: 2212–2228 e12

Zhang Q, Chen CZ, Swaroop M, Xu M, Wang L, Lee J, Wang AQ, Pradhan M, Hagen N, Chen L, Shen M, Luo Z, Xu X, Xu Y, Huang W, Zheng W, Ye Y (2020) Heparan sulfate assists SARS-CoV-2 in cell entry and can be targeted by approved drugs in vitro. Cell Discov 6: 80

Zhang Y, Wang S, Wu Y, Hou W, Yuan L, Shen C, Wang J, Ye J, Zheng Q, Ma J, Xu J, Wei M, Li Z, Nian S, Xiong H, Zhang L, Shi Y, Fu B, Cao J, Yang C et al. (2021) Virus-Free and Live-Cell Visualizing SARS-CoV-2 Cell Entry for Studies of Neutralizing Antibodies and Compound Inhibitors. Small Methods 5: 2001031

Zhao J, Guo S, Yi D, Li Q, Ma L, Zhang Y, Wang J, Li X, Guo F, Lin R, Liang C, Liu Z, Cen S (2021) A cell-based assay to discover inhibitors of SARS-CoV-2 RNA dependent RNA polymerase. Antiviral research 190: 105078

Zhong J, Gastaminza P, Cheng G, Kapadia S, Kato T, Burton DR, Wieland SF, Uprichard SL, Wakita T, Chisari FV (2005) Robust hepatitis C virus infection in vitro. Proceedings of the National Academy of Sciences of the United States of America 102: 9294–9

